# Task-Dependent Modulation of Feedback Control in Human Steering

**DOI:** 10.64898/2026.05.27.728136

**Authors:** Jo-Yu Liu, James R.H. Cooke, Luc P.J. Selen, W. Pieter Medendorp

**Affiliations:** Radboud University, Donders Institute for Brain, Cognition and Behaviour, Nijmegen, The Netherlands

**Author notes:** **Corresponding author:** Prof. dr. W.P. Medendorp, Donders Institute for Brain, Cognition and Behaviour, Radboud University, The Netherlands.

**Keywords:** steering, perturbations, optimal feedback control, task-specificity

## Abstract

We examined whether human steering behavior conforms to optimal feedback control (OFC) principles when driving a vehicle through sequences of upcoming gates varying in width (narrow/wide) relative to the vehicle’s size, while occasional lateral velocity perturbations elicited corrective steering responses. In 24 participants, three predictions of OFC were tested: (1) greater positional variability when passing wide gates; (2) reduced corrective steering (lower feedback gains) to perturbations for wide compared to narrow gates (minimum intervention); and (3) modulation of feedback gains according to the relative spatial configuration of upcoming gates. Steering data were analyzed in terms of vehicle position and control velocity, the latter representative of feedback gain modulations. Results supported all predictions. When no perturbations occurred, participants showed wider distributions of passing locations through wide gates, consistent with allowing more task-relevant variability. During perturbed trials, feedback gains were lower for wide gates, confirming context-dependent control. When both gate types were center aligned, feedback responses were symmetric, whereas alignment of inner posts produced asymmetric feedback gains, being stronger for outward perturbations in narrow gates. Overall, these findings indicate that participants adapt their feedback control strategies to task-specific spatial constraints, demonstrating flexible optimization of steering control behavior.

## Introduction

Driving a vehicle through gates of different widths requires the central nervous system (CNS) to integrate sensory feedback with predictive motor commands (Jansen and Fajen 2024, 2025; Tuhkanen et al. 2019). In such a scenario, steering must be regulated based on visual cues, like gate width and distance, together with non-visual cues about self-motion (Land and Horwood 1995; Markkula et al. 2023; Salvucci and Gray 2004). When the gate is narrow, fine steering adjustments are needed to minimize collision risk, while steering through wider gates can be achieved with coarser control (Benderius and Markkula 2014; van der El et al. 2023; Haselberger et al. 2025; Land and Horwood 1995; Lappi and Mole 2018; Mecheri et al. 2017). What theoretical principles underpin these observations?

A prominent theory of human motor control is optimal feedback control (OFC, (Scott 2012; Todorov and Jordan 2002), which models the CNS as a stochastic controller whose feedback gains minimize an expected cost function comprising task accuracy and control effort. A central prediction of this framework – the minimum intervention principle (MIP) – proposes that the CNS corrects only movement deviations that affect task performance while ignoring task-irrelevant variability. Empirical evidence for this principle has been found across eye movement, reaching and object avoidance tasks (Chen-Harris et al. 2008; Keyser et al. 2017; Liu and Todorov 2007; Nashed et al. 2012). Overall, feedback gains are thought to adjust dynamically in response to factors such as the behavioral goal (Nashed et al. 2012), time-to-target (Česonis and Franklin 2020; Dimitriou et al. 2013), the direction of movement deviations (Franklin et al. 2017), and the stability of the environment (Franklin and Wolpert 2008).

Recent studies have extended OFC theory to driving and steering behavior (Liu et al. 2025; Nash and Cole 2020). In this case, the CNS constructs and updates an internal model that maps the steering commands onto resulting changes in vehicle state (van Helvert et al. 2025; Lappi and Mole 2018; Liu et al. 2025). This internal model enables the selection of a movement policy that achieves the steering goal (e.g., steer the vehicle through a gate) and minimizes control effort (e.g., avoid unnecessary steering actions). The controller is defined as a set of state dependent feedback gains that define the strength of the corrective responses. This controller is driven based on the vehicle’s state (position and velocity) estimate that is derived from sensory information (visual, vestibular, proprioceptive) and prior information, like earlier issued control signals.

Feedback gains can experimentally be assessed by introducing controlled perturbations and measuring compensatory responses. In a recent study, employing a self-steerable motion platform, we investigated how participants steered a vehicle along roads of varying widths while subjected to continuous, time-varying mechanical perturbations (Liu et al. date unknown). Only limited support for the minimum intervention principle was found; steering feedback gains remained largely invariant across road widths. One possible explanation is that the control policy was tuned for maintaining the vehicle on the road within acceptable limits, trading efficiency for robustness (Başar and Bernhard 1995; Crevecoeur et al. 2019; Maurus et al. 2023).

To further investigate the minimum intervention principle in steering, the present study used a more standard perturbation paradigm, examining how steering feedback gains consider the relevance of a single perturbation for task performance. Specifically, we measured feedback gains in response to short-lived mechanical perturbations delivered immediately before participants drove through an upcoming narrow or wide gate (see Fig. 1B). We compared how participants changed their steering velocity when perturbed in the direction of the gate (outward perturbation) versus away from the gate (inward perturbation) under center-aligned or inner-post-aligned gate conditions (Fig 1D).

**Figure 1.**
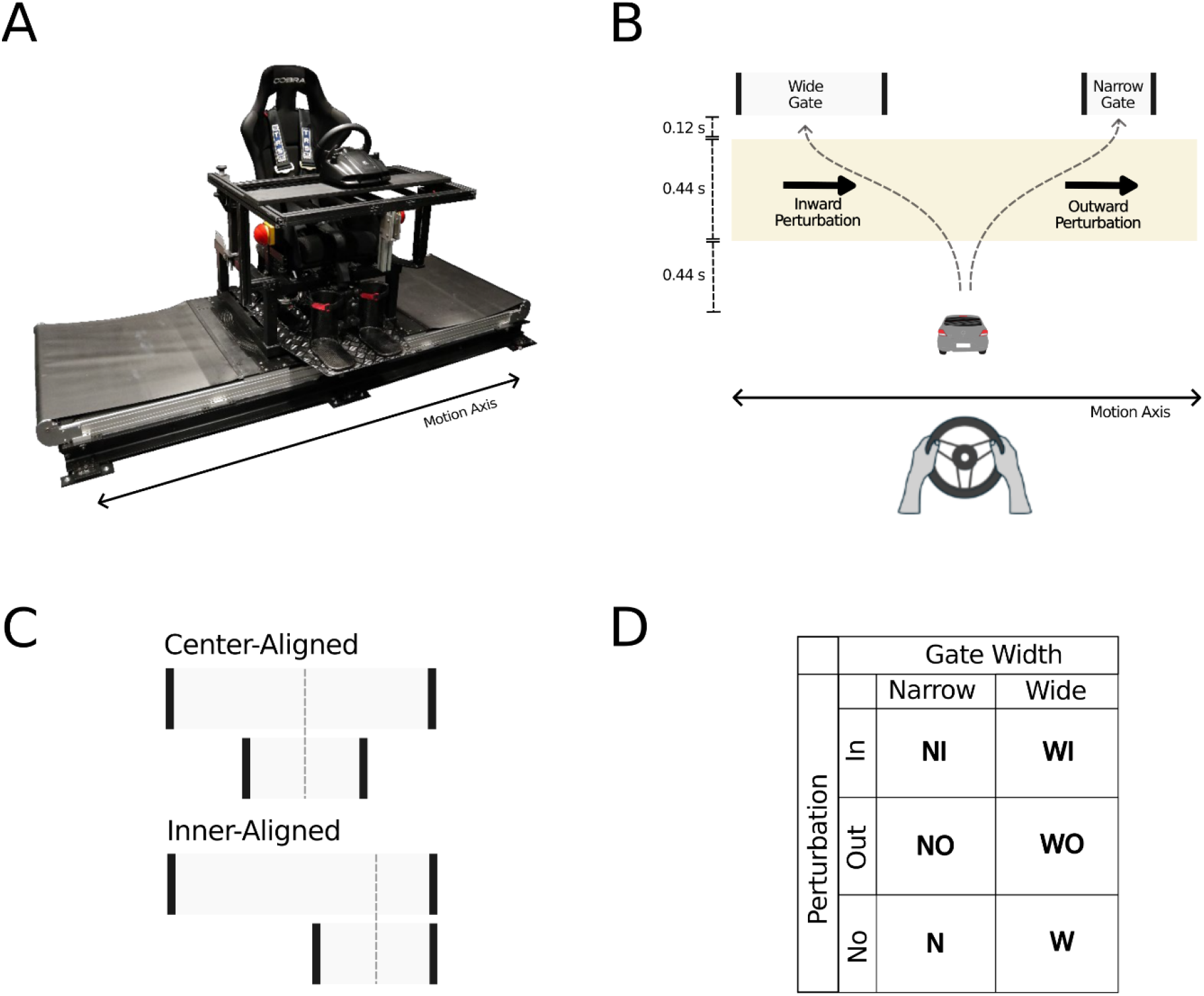
Experimental setup and paradigm. A. Linear motion platform (“vestibular sled”). B. Task: participants steered a cartoon vehicle whose on-screen lateral motion matched the sled’s movement, passing it through narrow or wide gates, either on the left or right. Occasionally, the sled and vehicle were perturbed outward (toward the gate) or inward (away from it) prior to gate passing. C. Twelve participants completed the center-aligned gate condition (both gate types centered), and twelve performed the inner-aligned condition (gate inner posts aligned, here shown for left-sided gates). D. Experimental design, showing all combinations of perturbation type (inward, outward, none) and gate width (narrow, wide).

When participants aim to steer toward the center of the gate, and the narrow and wide gates are aligned by their centers, both inward and outward deviations are expected to elicit higher feedback gains for the narrow gate than for the wide one, reflecting the narrower gate’s smaller steering margins. In contrast, if drivers aim for trajectories that pass near the gate’s inner post, another prediction of OFC, then when the narrow and wide gates are aligned by their inner posts, inward perturbations should produce similar feedback gains because both gates allow comparable passing space. In contrast, outward perturbations should trigger a higher gain for the narrow gate than for the wide one, reflecting their differences in passing space (Land and Horwood 1995; Salvucci and Gray 2004).

## Methods

### Participants

Twenty-four naïve participants (7 male, 17 female), ranging in age from 18 to 42 years, without any history of motion sickness, took part in the study. The study was approved by the ethics committee of the Faculty of Social Sciences of Radboud University Nijmegen (no. ECSW-2022-082), the Netherlands. All participants gave written informed consent. The experiment lasted approximately 60 minutes per participant including instructions and administration. Participants were compensated with course credits, or €15.00.

### Setup

Participants were seated in a chair on top of a custom-built linear motion platform (called the vestibular sled, Figure 1A) that could be moved passively by the experiment software or actively by the participant using a steering wheel, or a combination of both. The participants’ head was restrained by two head clamps, and their body was secured by a five-point seat belt. Their lower legs and feet were also strapped. The sled moved along a track (0.9 m long) driven by a linear motor (TB15N; Tecnotion, Almelo, The Netherlands) and controlled by a servo drive (Kollmorgen S700; Danaher, Washington, DC). The motion axis of the platform was aligned with the participant’s interaural axis. Emergency buttons on either side of the sled could be pressed to stop the sled motion at any time. A steering wheel (Logitech G27 Racing Wheel, Lausanne, Switzerland) was mounted at a comfortable handling distance, approximately 0.50 m in front of the participants’ chest. Participants placed their hands on the steering wheel at the 9 and 3 o’clock positions. The wheel’s orientation, which was sampled at 100 Hz, had a range between −450° and +450° and a resolution of 0.06°. The steering wheel angle was multiplied by 0.008 m/s/° to determine the instantaneous velocity of the sled, with the 12 o’clock direction defined as 0 m/s. Thus, a 45° rotation of the steering wheel was equivalent to a lateral sled velocity of 0.36 m/s. Sled velocity was capped at 1 m/s. Occasionally, the sled was subjected to a mechanical perturbation following a Gaussian velocity profile, characterized by a standard deviation of 0.075 s, which peaked 0.33 s before gate crossing with a peak velocity of 0.6 m/s. This short-lived perturbation would introduce a sled displacement of 11.2 cm, if uncompensated by the participant.

Participants viewed a 77-inch OLED screen (OLED77C3PUA; LG, Seoul, South Korea) with a resolution of 1920×1080 pixels and a refresh rate of 120 Hz, positioned centrally in front of the participant at 1.6 m. The bottom of the display showed a cartoon-style vehicle, with a width of 0.084 m, whose lateral position matched that of the participant on the sled. Gates were depicted as two vertical white lines (10 cm tall), separated by a fixed distance. Gates were either narrow (0.15 m width) or wide (0.4 m width). A gate appeared at the top of the screen, offset left or right relative to the car, and descended downward at a constant speed of 0.75 m/s for a duration of 1 s (Fig 1B).

The experiment was controlled by custom code written in Python 3 using the PsychoPy module v.3.6.9 (Peirce 2008).

### Behavioral task

Using the steering wheel, the participants’ task was to control the sled’s lateral motion to ensure the on-screen vehicle passed through an approaching gate. Occasionally, the sled’s motion was briefly perturbed by a velocity pulse, peaking 0.33 s before passing the upcoming gate. The pulse was either *outward* (toward the gate, with the vehicle motion) or *inward* (away from the gate, against the vehicle motion) (Figure 1B). For short, we refer to these as perturbed gates. When a gate was passed successfully (defined as the entirety of the car passing between the two lines that represent gate posts), the car flashed green for 1 s; if unsuccessful, it flashed red. The duration for moving from one gate to the other was 1 s. When passing a gate at the bottom of the screen, a new gate would immediately appear at the top of the screen.

Subsequent gates alternated sides on the screen to keep the sled within its motion range of 0.9 m. All perturbed gates were spawned at 22.5 cm or 35 cm (depending on condition, see below) relative to the participant’s position when passing the prior gate. All unperturbed gates were spawned at fixed positions on the screen, either 11.15 cm or 17.5 cm left or right of the screen’s center. This minimized drift toward the track edges. A run consisted of 64 consecutive gates, producing a slalom-like movement of the sled and vehicle. Of these 64 gates, 16 were perturbed gates, which were equally distributed across gate side (left/right), gate width (narrow/wide) and perturbation direction (inward/outward) (see Fig 1D). A perturbed gate was always followed by an unperturbed gate. Each participant completed a total of 20 runs, each lasting 76 s (including a 3 s countdown at the start, and a 6 s grace period before the end of the run). There was a short break after every run.

Before the actual experiment began, participants performed first a practice run with only unperturbed gates to familiarize themselves with the mapping between the steering wheel motion, the physical sled motion and the visual vehicle motion on the screen.

Next, participants completed two practice runs that included the same perturbations as in the actual experimental runs. In total, a participant completed 23 runs, of which the final 20 were the experimental runs. These experimental runs resulted in 320 perturbed and 960 unperturbed gates. For each combination of gate side (left/right), perturbation direction (inward/outward) and gate width (narrow/wide), this resulted in 40 perturbed trials.

## Conditions

The 24 participants were randomly assigned to two groups of 12. One group was tested in the *center-aligned* gate condition (Fig 1C), in which both the centers of both the narrow and wide perturbed gates were presented with lateral offsets of 0.35 m relative to the passing location of the previous gate. In this case, both inward and outward perturbations are expected to produce higher feedback gains for the narrow than for the wide gate, reflecting the narrow gate’s reduced steering margins. The second group was tested in the inner-post-aligned gate condition (*inner aligned*, Fig 1C). Here, the narrow and wide perturbed gates had center offsets of 0.225 and 0.35 m, respectively, relative to the passing location of the previous gate. According to OFC predictions, inward perturbations should evoke similar feedback gains due to comparable passing tolerances, while outward perturbations should produce higher gains for the narrow than the wide gate.

### Data analysis

Data were processed offline via Python 3, using NumPy (Harris et al. 2020), SciPy (Virtanen et al. 2020), and statsmodel (Seabold and Perktold 2010) modules. The first three runs from each participant were practice runs and excluded from analysis. We then split each run into 64 trials, each containing the 1 s steering wheel angles and sled/vehicle motion trajectories to a specific gate. The steering wheel angle was converted into sled velocity. On perturbed gates, the sled motion contained an additional velocity pulse. In total, each participant performed 960 unperturbed and 320 perturbed trials. Of the perturbation trials, 160 had an inward perturbation, and 160 had an outward perturbation (Figure 1D). Half of these perturbations were for narrow and half for wide gates.

The success rate for the vehicle passing through the gate was defined in binary terms. A trial was considered successful if the entire vehicle was within the two gate posts when passing. If any part of the vehicle overlapped with a gate post while passing through, it was classified as a miss.

We examined whether the motion trajectories to the wide and small gate differed in the presence and absence of a perturbation. We specifically focused on differences in the passing location distribution of the vehicle between wide and narrow gates. To obtain the passing locations, we aligned all trajectories on the center of the gate location and added the standard gate center distance (so 22.5 cm and 35 cm).

We examined the corrective responses to the perturbation separately for each gate type by comparing steering wheel excursions relative to their most similar unperturbed baseline trial. The set of possible baseline trials only comprised trials in which there had been no perturbation towards the starting gate and finishing gate. Similarity between two trajectories was quantified as the reciprocal of the root mean squared (RMS) error of their perpendicular deviation; that is, a smaller RMS error indicated greater similarity. To minimize contamination from residual effects of the previous trial and early perturbation influences, we restricted the similarity calculation to the 350–650 ms segment of the trajectory (see Keyser et al., 2017, for a comparable method). The unperturbed trial that yielded the lowest RMS error was taken as the best matching baseline.

### Statistics

Analyses were performed separately for the center-aligned and inner-aligned experimental groups. Statistical tests were evaluated against a two-tailed alternative hypothesis, unless noted otherwise. Results were considered significant at an alpha level of 0.05, unless mentioned otherwise.

To analyze success rates, we used generalized linear models for binomial data and a logit link function. The fixed factors were gate width (narrow vs. wide), perturbation direction (inward, none, outward), and gate side (left vs. right), with participant included as a random effect.

The mean gate-passing location and their standard deviation were first analyzed using a three-way repeated-measures ANOVA, with factors gate width (narrow/wide), perturbation direction (inward/none/outward) and gate side (left/right). As expected, gate side did not affect any of the outcome measures. Therefore, we present results from two-way repeated-measures ANOVAs where we adjusted for and collapsed over gate sides.

To assess the corrective steering responses, relative to the unperturbed steering wheel trajectories, we performed running, repeated-measures ANOVAs with perturbation direction (inward/outward), gate width (narrow/wide) as within-subject factors. A running test was considered significant once it remained continuously significant at a conservative P<10^−5^ level until the end of the trial. This approach enabled us to pinpoint the time intervals when conditions began to differ significantly while mitigating the multiple comparison problem. Cluster-based permutation tests gave similar results (not reported).

The baseline corrected control velocity value at gate passing was taken as a proxy for the feedback gain and was analyzed using a repeated-measures ANOVA with perturbation direction and gate width as main factors.

### Data availability

Upon publication, all data will be made publicly available at the Radboud Data Repository: *https://data.ru.nl/*, via a persistent identifier reserved for this collection.

## Results

Our experiment was designed to test whether physical perturbations during gate-directed steering result in gate width-dependent corrections of the steering trajectories. Participants steered either toward an upcoming narrow or a wide gate. In 25% of the trials, unknown to the participant, the sled was perturbed by a transient velocity pulse, peaking 330 ms before passing the upcoming gate, directed in either outward or inward direction relative to the gate. According to OFC, it is expected that the spread of passing locations in unperturbed trials is larger for the wide compared to the narrow gates, due to a reduction in corrective responses to fluctuations caused by internal noise. Similarly, smaller corrective responses are expected for perturbed trials in which the participant was steering towards a wide gate and most likely these responses also show larger spread compared to a narrow gate.

Figure 2A illustrates the vehicle’s lateral position over time, combined across left and right gates (flipping the data for the left gate) and averaged across the 12 participants, for the center-aligned condition. Trajectories are aligned on the center location of the wide gate. The inset displays the spatial trajectories separately for left and right gates. Dashed lines indicate trajectories toward wide gates, while solid lines correspond to those toward narrow gates. Gray traces represent unperturbed trials, and green and orange traces depict outward and inward perturbed trials, respectively. Note that perturbed trajectories start at a fixed location relative to the upcoming gate, whereas unperturbed trajectories do not (their gates spawned at fixed locations in space).

**Figure 2.**
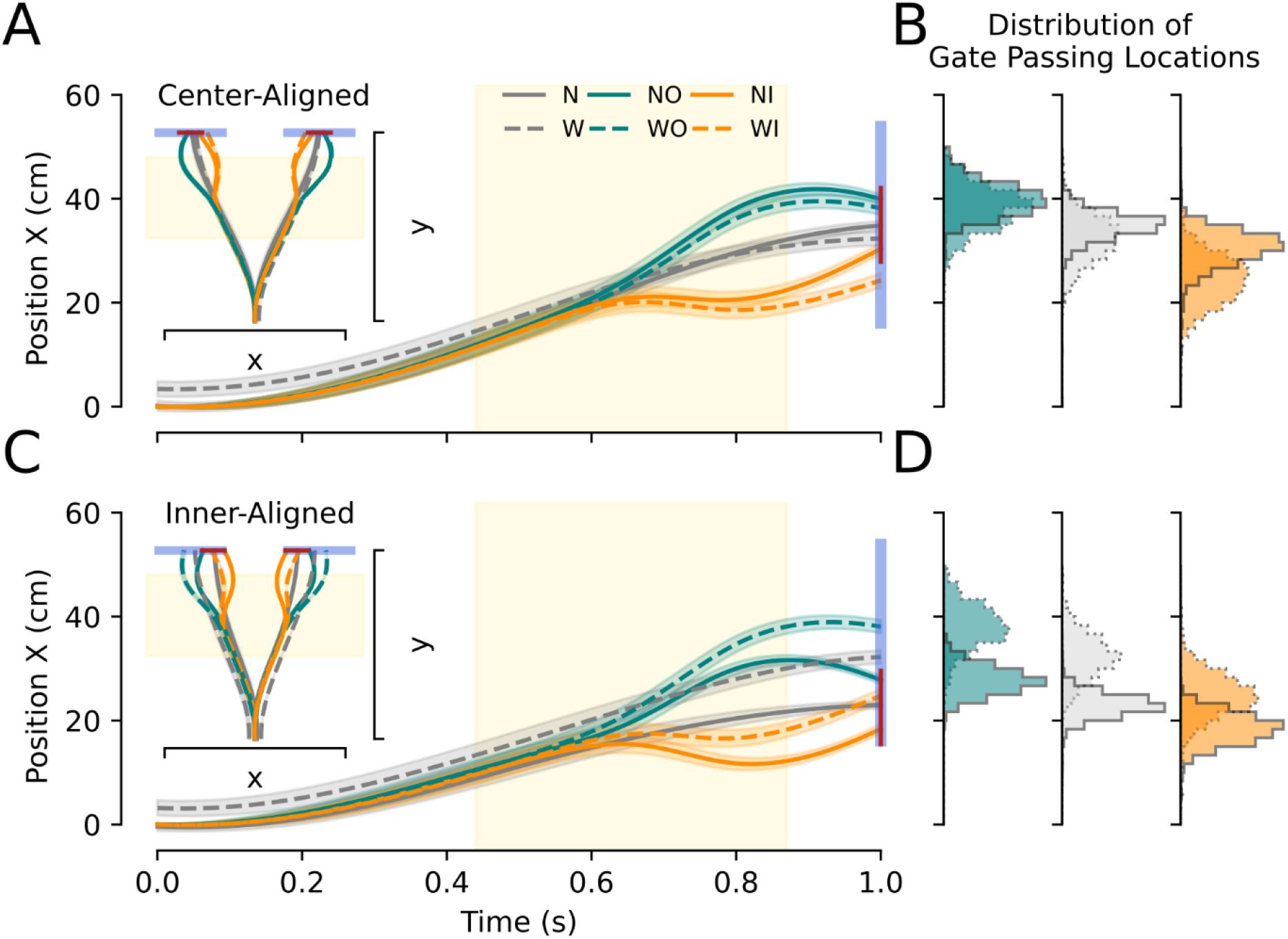
Motion trajectories towards the gate and distribution of gate passing location. A. Mean position trajectories across participants in the center-aligned condition. Blue and thin red bars mark wide and narrow gates, respectively. Solid lines indicate narrow-gate trajectories; dashed lines indicate wide-gate trajectories. Gray represents no perturbation, green outward, and orange inward perturbations. Inset displays spatial trajectories, separately for left and right gates. B. Distributions of gate passing locations; same color and style conventions. C. Mean position trajectories across participants in the inner-aligned condition. Format as in A. D. Distributions of gate passing locations, format as in B.

In the unperturbed trials, the mean sled trajectories for the narrow gate pass through the gate center, but for wide gates they show a slight deviation toward the inner post (Fig. 2A). This is in accordance with OFC, which emphasizes trading-off effort and success. That is, for wide gates, participants can reduce effort by cutting the corner while maintaining a high likelihood of successful passage. This also explains why trajectories toward the wide gate already start from an offset position: the offset arises from cutting the corner of the previous wide gate on the opposite side.

Following perturbation, the sled trajectories for the inward and outward perturbations began to diverge at about 650 ms (Fig 2A) but converge when approaching the gate. For inward perturbations, corrective movements were stronger for narrow than wide gates, as needed to successfully pass the gate. For outward perturbations the corrections look very similar for the narrow and wide gate. A repeated-measures ANOVA, including unperturbed trials, revealed a significant main effect of gate width on the passing locations (narrow: 35.05 cm, (4.18 SD), wide: 31.62 cm (6.30 SD); F(1,11)=38.02, p=0.0001, np^2^=0.77). In addition, there was a significant effect of perturbation (none, inward, outward) (F(2,22)=256.38, p<10^−4^, np^2^=0.96) and a significant interaction between perturbation and gate width (F(2,22) = 107.58, p < 10^−4^, np^2^ = 0.90). Post-hoc paired t-tests show significant differences in passing locations between narrow and wide targets (Fig 2B) for the unperturbed (t(11)=4.58, p<0.001), outward-perturbed (t(11)=9.80, p<0.001) and inward-perturbed trials (t(11)=2.92, p=0.01).

Fig 2B illustrates the distributions of passing locations, showing greater spread for wide than narrow gates, irrespective of the presence or absence of perturbations. This is in line with OFC. We performed a two-way repeated measures ANOVA with main factors gate width (narrow, wide) and perturbation (none, inward, outward) to examine their effects on the standard deviation of the passing locations. Indeed, there was a significant main effect of gate width (F(1,11)=80.55, p<10^−5^, np^2^ = 0.88) but not of perturbation (F(2,22)=0.05, p=0.95, np^2^ = 0.01), confirming the overall smaller spread of passing locations for narrow gates. In addition, there was a small interaction effect (F(2,22)=5.68, p=0.01, np^2^=0.51). Post-hoc paired samples t-tests indicate that this gate width dependence is mainly observed in the unperturbed and inward perturbed trials (t(1,11)=11.45, p<0.001 and t(1,11)=4.61, p<0.001, respectively), while for outward perturbations the effect did not reach statistical significance (t(1,11)=2.01, p=0.06).

Although the results obtained so far provide general support for the principles of OFC theory, we tested an additional group of 12 participants under the inner-aligned condition to further evaluate the theory. Based on OFC predictions, steering trajectories should pass closer to the gate’s inner post if accuracy constraints allow so, as indeed seen in Fig. 2A and C for the wide gates. Therefore, we reasoned that when the narrow and wide gates are aligned by their inner posts, inward perturbations should produce similar corrections, given more similar passing space. In contrast, outward perturbations should yield a stronger correction for the narrow than wide gate than for the wide one, reflecting their differing passing space.

Figures 2C and 2D show sled trajectories for inner-aligned gate locations, in the same format as Figures 2A and 2B. The unperturbed trajectory for the wide gate shows a bias towards the inner post but does not go as close as the trajectory to the narrow gate. Nevertheless, following inward perturbations, the trajectories exhibit more similar corrective patterns across both gate widths than those following outward perturbations (Fig. 2C). Specifically, outward perturbations evoke stronger corrections for the narrow gate, whereas towards the wide gate only modest compensation is seen. This pattern is also reflected in the distribution of passing locations for the two gate widths, which show more overlap for inward than outward perturbations (Fig. 2D). Indeed, quantitatively, for inward perturbations, the mean passing locations are 24.8 and 18.4 cm (difference = ~6 cm) for the wide and narrow gate, respectively, while for outward perturbations they are 38.07 and 27.79 cm (difference = ~10 cm). These observations are confirmed in a repeated-measures ANOVA, showing significant main effects of gate width (F(1,11)=562.81, p<10^−5^, np^2^=0.99) and of perturbation (F(2,22) = 295.61, p< 10^−5^, np^2^ = 0.96) on the passing locations. We also found a significant interaction effect between perturbation and gate width (F(2,22)=64.70, p<10^−5^, np^2^=0.85). The passing location distributions show smaller standard deviations for narrow than wide gates for both inward (narrow: 2.66 cm (0.98 SD), wide: 3.98 cm (1.13 SD)) and outward perturbations (narrow: 2.42 cm (0.49 SD), wide: 3.99 cm (0.72 SD)). This width effect was statistically significant (F(1,11)=210.77, p<10^−5^, np^2^=0.93) and interacted with the perturbation condition (F(2,22)=8.35, p=0.002, np^2^ = 0.37).

While Fig. 2 illustrates that steering trajectories generally passed through the gate following perturbations, there was some variability in individual participants’ success rates. Fig. 3 displays these success rates, combined across leftward and rightward gates, for each participant (small dots) and the group’s mean (solid and dashed circles for the narrow and wide gates, respectively). For unperturbed trials, success rate was very high (p(success) > 0.980) across both narrow and wide gates. For perturbed trials, success rate was more variable across participants (0.36 - 1.00) and lower for narrow (0.36 - 0.98) compared to wide (0.83 - 1.00) gates. This difference was significant in both the center-aligned (χ^2^(1, N = 72) = 27.37, p < 0.001) and the inner-aligned gate locations (χ^2^(1, N = 72) = 36.77, p < 0.001).

**Figure 3.**
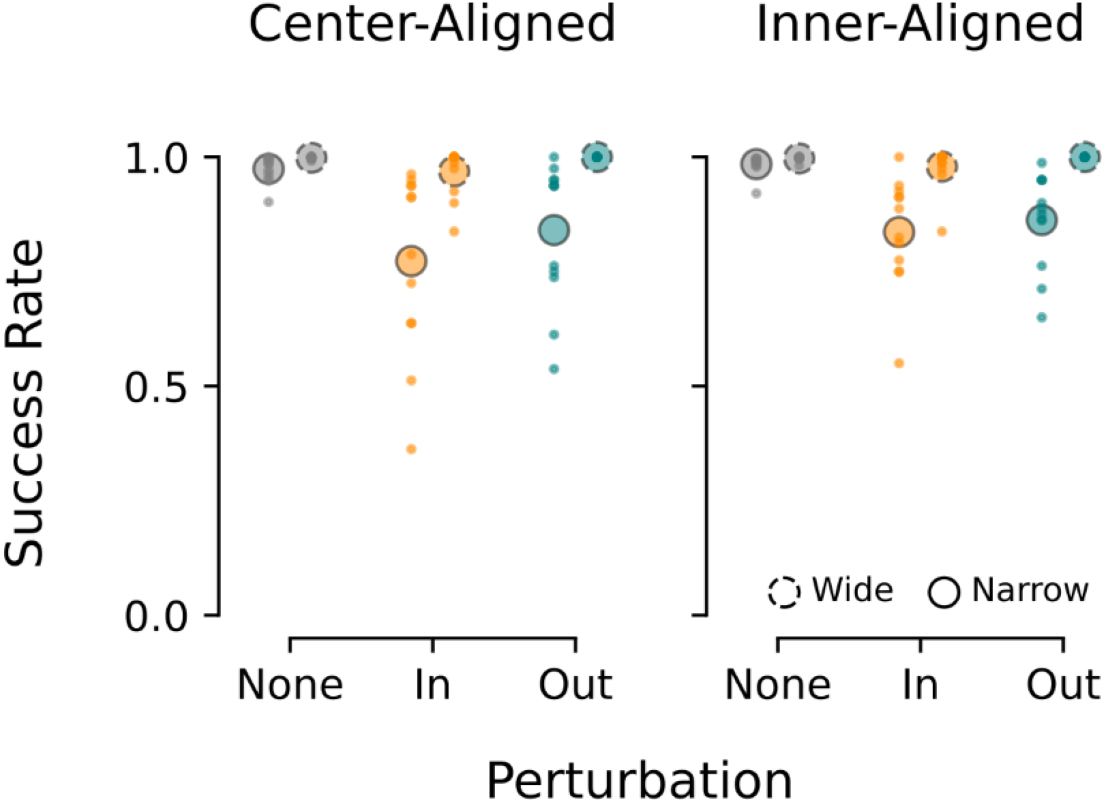
Success rates in gate passing in the center- and inner-aligned gate conditions. Small dots, individual participants. Large circles, group means.

For a closer examination of the change in reactive control to perturbations over time, Fig. 4 plots the changes in lateral steering control velocity by the participants, separately for each gate width and perturbation direction. These velocity traces follow from comparing steering wheel excursions relative to their most similar unperturbed baseline (see Methods).

**Figure 4.**
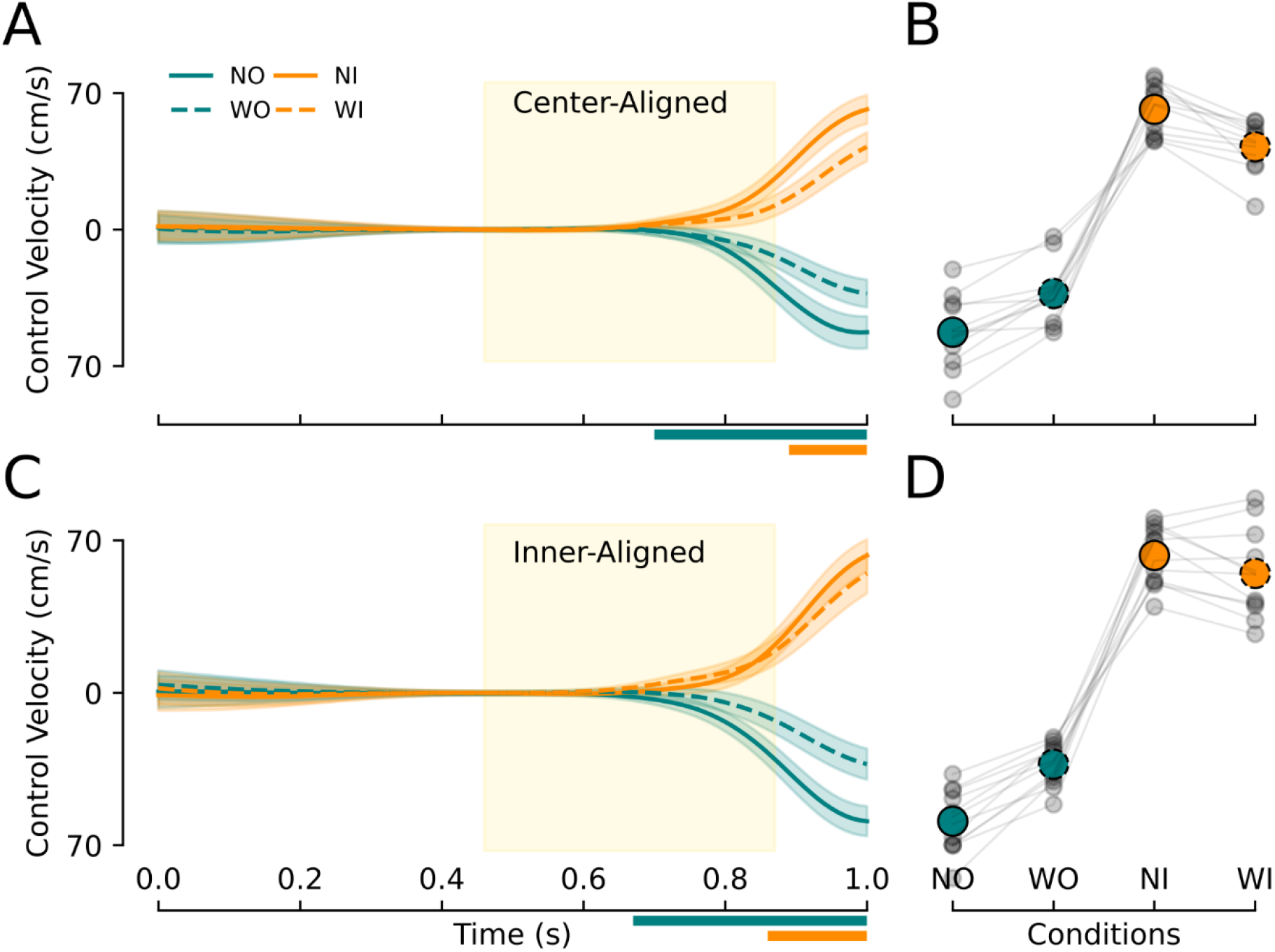
Steering control velocity as a function of time. A. Mean steering wheel velocity relative to matched unperturbed trajectories in the center-aligned condition. Solid and dashed traces denote narrow and wide gates, respectively. Green trace: outward perturbation; orange: inward perturbation. Colored bars indicate time points of significant factors: green = perturbation direction, orange = gate width. B. Control velocity at gate passing, with individual participant data in gray. C. Same as A for the inner-aligned condition. D. Same as B for the inner-aligned condition.

In the center-aligned condition (Fig. 4A), consistent with the positional data shown in Figure 2A, the mean responses across participants show that both inward and outward perturbations elicited stronger control velocities for the narrow gate compared to the wide gate. Note that the participant is still controlling the sled motion while passing through the gate (i.e., velocity is not zero). This pattern is consistent with OFC. A significant effect of perturbation direction, i.e. inward being dissociable from outward, emerged at 0.70 s, i.e., just after peak perturbation velocity (Fig. 4A horizontal green bar). Differences in control velocities due to gate width reached significance at 0.89 s (i.e., 0.22 s after perturbation peak (Fig 4A. orange bar). Fig 4B further illustrates the individual participants’ control velocities when passing through the gate, for each participant (gray circles) and the group’s average (filled and open circles for the narrow and wide gates, respectively). A repeated-measures ANOVA, where we flipped the sign of the outward perturbations to be able to compare magnitudes, showed a significant main effect of gate width (F(1,11)=157.74, p<10^−6^, ηp^2^=0.93), but neither perturbation direction (F(1,11)=12.20, p=0.005, ηp^2^=0.59) nor their interaction reached significance (F(1,11)=6.86, p=0.024, ηp^2^=0.38).

For the inner-aligned condition, based on Fig 2C, we expected similar perturbation responses for the narrow and wide gate, while outward perturbations should evoke stronger changes in control velocity for the narrow compared to the wide gate. This pattern is confirmed in Fig 4C. Again, inward and outward perturbations became dissociable around 0.67 s (i.e., at perturbation peak, Fig. 4C green bar) and the effect of gate width reached significance at 0.86 s, i.e., 0.19 s after perturbation peak (Fig. 4C orange bar). Fig 4D summarizes the control velocities at gate passing, for each participant (gray circles) and the group’s average (filled and open circles). Indeed, a repeated-measures ANOVA showed a significant main effect of gate width (F(1,11)=41.93, p<10^−4^, ηp^2^=0.79), but no other significant effects.

## Discussion

We investigated whether human participants’ steering behavior aligns with the principles of optimal feedback control (OFC) theory when they steered a car through a series of upcoming gates that were either wide or narrow relative to the vehicle’s size. In 25% of the trials, we introduced perturbations to the car’s lateral velocity to evoke corrective responses. Three predictions were made based on OFC theory: First, participants will allow greater natural variability in their steering when faced with a wide gate, resulting in a broader distribution of gate passing locations during unperturbed trials. Second, participants should exhibit less corrective steering, i.e. lower feedback gains, in response to perturbations compared to when steering through narrow gates (the minimum intervention principle). Finally, OFC also predicts that participants consider the entire steering scenario, adjusting their feedback gains according to the relative positions of the upcoming gates.

Our findings supported all these predictions. In the unperturbed trials, participants passed through wide gates more successfully (Fig 3), showing broader passing distributions compared to the narrow gates (Fig 2). This confirms the first prediction. In response to perturbations, feedback gains differed between narrow and wide gates, validating the second prediction. When both gate types were centered at the same position relative to the previous gate, feedback gains were similar for inward and outward perturbations (Fig 4B). However, when the inner posts of the gates were aligned, feedback gains became asymmetric: inward perturbations evoked comparable feedback gains for both gate widths, while outward perturbations elicited stronger feedback gains for the narrow than wide gates (Fig 4D). This asymmetry confirms the third prediction. These effects were observed not only in the position trajectories (Fig 2), but even more clearly in the modulation of steering control velocity (Fig 4).

Together, these findings indicate that participants adapted their feedback control policies to the gate-specific spatial constraints, consistent with the principles of OFC. Our results demonstrate that the sensorimotor system for steering flexibly regulates feedback responses to balance task goals and control costs, optimizing control in relation to both immediate and future environmental constraints (Liu et al. 2025). These task-dependent constraints link feedback gains to gate width and placement of future gates. Especially, the latter could account for the distinct patterns of feedback gain observed between center- and inner-aligned conditions. This suggests that the CNS adapts to the statistical properties of the environment when such variations are behaviorally relevant. In particular, we observed asymmetric gains for the wide gate in the inner aligned condition, with a higher gain for the inward compared to the outward perturbation. Such direction-dependent feedback gains have also been observed during reaching, where participants upregulated their visuomotor feedback gains under task-relevant directional perturbations and lowered their gain transient task-irrelevant perturbations (Franklin and Wolpert, 2008). Most importantly, they also show that these adaptations can be direction specific (Franklin et al. 2014) exp. 2).

Our finding that steering feedback gains varied with gate width indicates that sensory inputs directly inform the steering controller. In our task, visual and vestibular signals detect perturbations and update the estimated vehicle state, i.e., its position and velocity relative to the gate. The controller applies feedback gains to this estimate, modulating the corrective steering responses. The dependence of these responses on gate width contradicts the idea that vestibular-driven corrections simply stabilize the vehicle in space (Blouin et al. 2015). Instead, their attenuation under wide-gate conditions, where steering demands are lower, supports a control strategy consistent with OFC (Todorov and Jordan 2002).

Steering velocity corrections began approximately 190-220 ms after the perturbation peak (see Fig 4). Under the assumption that the effective perturbation starts 3SDs earlier (~20 ms), we find a value of 210-240 ms, which is consistent with latencies reported in previous studies examining vestibular evoked feedback responses in reaching (Keyser et al. 2017; Moreau-Debord et al. 2014; Oostwoud Wijdenes et al. 2019; Smith and Reynolds 2017). Moreover, in the center-aligned condition, the ~40% reduction in feedback gain in the wide gate condition closely matches findings of 35% to 45% reduction for visual and proprioceptive perturbations in reaching tasks, respectively (Keyser et al. 2019; Knill et al. 2011). Our results also align with the 33% reduction reported for vestibular-induced perturbation corrections for wide compared to narrow target reaches obtained through galvanic vestibular stimulation (Keyser et al. 2017).

One might ask why corrections toward the wide gate were not entirely suppressed, given that the wide gate would also be passed without them. After all, the induced displacement is only 11.2 cm, which is much smaller than the gate’s 20 cm half width. First, it is plausible that the modulation of feedback gain, which depends on gate width, overlaps temporally or functionally with other feedback responses that occur independently of gate width (Keyser et al. 2017; Nashed et al. 2012). Second, from an optimal feedback control perspective, a controller balancing accuracy and effort would still correct lateral perturbations to pass the gate with zero lateral velocity (Liu and Todorov 2007). Delaying these corrections would demand larger steering adjustments later. Thus, the optimal strategy maintains a reduced, but nonzero feedback gain even when navigating wide gates (Franklin and Wolpert 2011). Third, zero feedback gain implies that participants could predict the perturbation and its effects on self-motion. However, since the perturbation was consistent in timing and duration, but randomized in direction and magnitude, this seems unlikely, suggesting they used a more cautious strategy to ensure successful crossing of gates. Finally, it remains possible that participants anticipated the whole sequence of gates, rather than treating each gate individually, and adjusted their feedback gains to optimize steering performance across the entire steering scenario.

What computations underlie the tuning of feedback gains? In OFC, internal models predict sensory inputs and generate motor outputs using inverse dynamics (Todorov and Jordan 2002). Because vestibular signals encode head acceleration, effective control of the hands on the steering wheel requires these signals to be integrated with neck proprioceptive input and transformed into a trunk-centered reference frame (Bakker et al. 2015, 2019; Mergner et al. 2001; Zobeiri and Cullen 2022). Based on the resulting estimate of self-motion, an internal inverse model of the biomechanics of the steering arms and interaction torques allows the brain to generate precise and task relevant motor commands. These internal models have been shown to be precise and flexible (Alefantis et al. 2022; Cullen 2023; van Helvert et al. 2022, 2025; Stavropoulos et al. 2022). For example, Moreau-Debord et al. (2014) showed that a complete model of arm biomechanics can successfully account for the observed magnitude of arm responses to galvanic vestibular stimulation. It is therefore not unreasonable to propose that the brain integrates self-motion control and steering within a unified feedback control policy.

Finally, we emphasize that the single-step perturbation paradigm used in this study is specifically designed to examine the principles of optimal feedback control. We do not suggest that this is the only control strategy that can be engaged during steering. In our earlier study, for instance, participants were instructed to keep a car within the road boundaries while continuous pseudorandom perturbations within a defined bandwidth were applied to the car’s lateral speed (Liu et al. date unknown). The results indicated that participants’ control strategies were tuned according to criteria that ensured the vehicle stayed within acceptable limits on the road, a finding more consistent with the principles of robust control theory (Başar and Bernhard 1995). Robust control focuses on controllers that maintain stability and satisfactory performance despite uncertainty in the model of the environment, as well as structure of variability and perturbations. This involves minimizing task error and control effect under a worst case scenario (Başar and Bernhard 1995; Crevecoeur et al. 2019; Maurus et al. 2023; Ueyama 2014). In contrast, OFC optimizes a feedback law to minimize expected cost under the assumption of an accurate internal model. These two frameworks, optimal feedback control and robust control, represent complementary control strategies that can coexist within the broader context of steering control. It may be an interesting avenue for future research to investigate the steering contexts that blur the line between optimal feedback and robust control.

## Acknowledgements

WPM, JYL and JRHC were supported by an NWA grant (NWA-ORC-1292.19.298). WPM is additionally supported NWO-SGW-406.21.GO.009 and Interreg NWE-RE:HOME.

## References

Alefantis P, Lakshminarasimhan K, Avila E, Noel J-P, Pitkow X, Angelaki DE. Sensory Evidence Accumulation Using Optic Flow in a Naturalistic Navigation Task. J Neurosci 42: 5451–5462, 2022.

Bakker RS, Selen LPJ, Medendorp WP. Stability of Phase Relationships While Coordinating Arm Reaches with Whole Body Motion. PloS One 10: e0146231, 2015.

Bakker RS, Selen LPJ, Medendorp WP. Transformation of vestibular signals for the decisions of hand choice during whole body motion. J Neurophysiol 121: 2392–2400, 2019.

Başar T, Bernhard P. H∞-Optimal Control and Related Minimax Design Problems: A Dynamic Game Approach. Birkhäuser Boston, 1995.

Benderius O, Markkula G. Evidence for a fundamental property of steering. Proc Hum Factors Ergon Soc Annu Meet 58: 884–888, 2014.

Blouin J, Bresciani J-P, Guillaud E, Simoneau M. Prediction in the Vestibular Control of Arm Movements. Multisensory Res 28: 487–505, 2015.

Česonis J, Franklin DW. Time-to-Target Simplifies Optimal Control of Visuomotor Feedback Responses. eNeuro 7: ENEURO.0514-19.2020, 2020.

Chen-Harris H, Joiner WM, Ethier V, Zee DS, Shadmehr R. Adaptive control of saccades via internal feedback. J Neurosci 28: 2804–2813, 2008.

Crevecoeur F, Scott SH, Cluff T. Robust Control in Human Reaching Movements: A Model-Free Strategy to Compensate for Unpredictable Disturbances. J Neurosci 39: 8135–8148, 2019.

Cullen KE. Internal models of self-motion: neural computations by the vestibular cerebellum. Trends Neurosci 46: 986–1002, 2023.

Dimitriou M, Wolpert DM, Franklin DW. The temporal evolution of feedback gains rapidly update to task demands. J Neurosci 33: 10898–10909, 2013.

van der El K, Pool DM, van Paassen MM, Mulder M. Modeling driver steering behavior in restricted-preview boundary-avoidance tasks. Transp Res Part F Traffic Psychol Behav 94: 362–378, 2023.

Franklin DW, Franklin S, Wolpert DM. Fractionation of the visuomotor feedback response to directions of movement and perturbation. J Neurophysiol 112: 2218–2233, 2014.

Franklin DW, Wolpert DM. Specificity of reflex adaptation for task-relevant variability. J Neurosci 28: 14165–14175, 2008.

Franklin DW, Wolpert DM. Computational Mechanisms of Sensorimotor Control. Neuron 72: 425–442, 2011.

Franklin S, Wolpert DM, Franklin DW. Rapid visuomotor feedback gains are tuned to the task dynamics. J Neurophysiol 118: 2711–2726, 2017.

Harris CR, Millman KJ, van der Walt SJ, Gommers R, Virtanen P, Cournapeau D, Wieser E, Taylor J, Berg S, Smith NJ, Kern R, Picus M, Hoyer S, van Kerkwijk MH, Brett M, Haldane A, del Río JF, Wiebe M, Peterson P, Gérard-Marchant P, Sheppard K, Reddy T, Weckesser W, Abbasi H, Gohlke C, Oliphant TE. Array programming with NumPy. Nature 585: 357–362, 2020.

Haselberger J, Böhle M, Schick B, Müller S. Exploring the Influence of Driving Context on Lateral Driving Style Preferences: A Simulator-Based Study. IEEE Trans Intell Transp Syst 26: 5448–5466, 2025.

van Helvert MJL, Selen LPJ, van Beers RJ, Medendorp WP. Predictive steering: integration of artificial motor signals in self-motion estimation. J Neurophysiol 128: 1395–1408, 2022.

van Helvert MJL, Selen LPJ, van Beers RJ, Medendorp WP. Internal models in active self-motion estimation: role of inertial sensory cues. J Neurophysiol 134: 171–182, 2025.

Jansen AJ, Fajen BR. Prospective control of steering through multiple waypoints. J Vis 24: 1, 2024.

Jansen AJ, Fajen BR. Visual control of steering through multiple waypoints. Sci Rep 15: 31261, 2025.

Keyser J, Medendorp WP, Selen LPJ. Task-dependent vestibular feedback responses in reaching. J Neurophysiol 118: 84–92, 2017.

Keyser J, Ramakers REFS, Medendorp WP, Selen LPJ. Task-dependent responses to muscle vibration during reaching. Eur J Neurosci 49: 1477–1490, 2019.

Knill DC, Bondada A, Chhabra M. Flexible, Task-Dependent Use of Sensory Feedback to Control Hand Movements. J Neurosci 31: 1219–1237, 2011.

Land M, Horwood J. Which parts of the road guide steering? Nature 377: 339–340, 1995.

Lappi O, Mole C. Visuomotor Control, Eye Movements, and Steering: A Unified Approach for Incorporating Feedback, Feedforward, and Internal Models. Psychol Bull 144: 981–1001, 2018.

Liu D, Todorov E. Evidence for the flexible sensorimotor strategies predicted by optimal feedback control. J Neurosci 27: 9354–9368, 2007.

Liu J-Y, Cooke J, Selen L, Medendorp P. Road width effects on feedback control in human steering behavior [Online]. date unknown https://www.authorea.com/users/1007478/articles/1367537-road-width-effects-on-feedback-control-in-human-steering-behavior [9 Apr. 2026].

Liu J-Y, Cooke JRH, Selen LJP, Medendorp WP. Visuoinertial and visual feedback in online steering control. PLoS Comput Biol 21: e1012659, 2025.

Markkula G, Lin Y-S, Srinivasan AR, Billington J, Leonetti M, Kalantari AH, Yang Y, Lee YM, Madigan R, Merat N. Explaining human interactions on the road by large-scale integration of computational psychological theory. PNAS Nexus 2: pgad163, 2023.

Maurus P, Jackson K, Cashaback JGA, Cluff T. The nervous system tunes sensorimotor gains when reaching in variable mechanical environments. iScience 26: 106756, 2023.

Mecheri S, Rosey F, Lobjois R. The effects of lane width, shoulder width, and road cross-sectional reallocation on drivers’ behavioral adaptations. Accid Anal Prev 104: 65–73, 2017.

Mergner T, Nasios G, Maurer C, Becker W. Visual object localisation in space: Interaction of retinal, eye position, vestibular and neck proprioceptive information. Exp Brain Res 141: 33–51, 2001.

Moreau-Debord I, Martin CZ, Landry M, Green AM. Evidence for a reference frame transformation of vestibular signal contributions to voluntary reaching. J Neurophysiol 111: 1903–1919, 2014.

Nash CJ, Cole DJ. Identification and validation of a driver steering control model incorporating human sensory dynamics. Veh Syst Dyn 58: 495–517, 2020.

Nashed JY, Crevecoeur F, Scott SH. Influence of the behavioral goal and environmental obstacles on rapid feedback responses. J Neurophysiol 108: 999–1009, 2012.

Oostwoud Wijdenes L, van Beers RJ, Medendorp WP. Vestibular modulation of visuomotor feedback gains in reaching. J Neurophysiol 122: 947–957, 2019.

Peirce JW. Generating stimuli for neuroscience using PsychoPy. Front Neuroinformatics 2: 10, 2008.

Salvucci DD, Gray R. A two-point visual control model of steering. Perception 33: 1233– 1248, 2004.

Scott SH. The computational and neural basis of voluntary motor control and planning. Trends Cogn Sci 16: 541–549, 2012.

Seabold S, Perktold J. Statsmodels: Econometric and Statistical Modeling with Python. Python in Science Conference, p. 92–96.

Smith CP, Reynolds RF. Vestibular feedback maintains reaching accuracy during body movement. J Physiol 595: 1339–1349, 2017.

Stavropoulos A, Lakshminarasimhan KJ, Laurens J, Pitkow X, Angelaki DE. Influence of sensory modality and control dynamics on human path integration. eLife 11: e63405, 2022.

Todorov E, Jordan MI. Optimal feedback control as a theory of motor coordination. Nat Neurosci 5: 1226–1235, 2002.

Tuhkanen S, Pekkanen J, Rinkkala P, Mole C, Wilkie RM, Lappi O. Humans Use Predictive Gaze Strategies to Target Waypoints for Steering. Sci Rep 9: 8344, 2019.

Ueyama Y. Mini-max feedback control as a computational theory of sensorimotor control in the presence of structural uncertainty. Front Comput Neurosci 8: 119, 2014.

Virtanen P, Gommers R, Oliphant TE, Haberland M, Reddy T, Cournapeau D, Burovski E, Peterson P, Weckesser W, Bright J, van der Walt SJ, Brett M, Wilson J, Millman KJ, Mayorov N, Nelson ARJ, Jones E, Kern R, Larson E, Carey CJ, Polat i, Feng Y, Moore EW, VanderPlas J, Laxalde D, Perktold J, Cimrman R, Henriksen I, Quintero EA, Harris CR, Archibald AM, Ribeiro AH, Pedregosa F, van Mulbregt P. SciPy 1.0: fundamental algorithms for scientific computing in Python. Nat Methods 17: 261–272, 2020.

Zobeiri OA, Cullen KE. Distinct representations of body and head motion are dynamically encoded by Purkinje cell populations in the macaque cerebellum. eLife 11: e75018, 2022.

